# Cooperative induction of ordered peptide and fatty acid aggregates

**DOI:** 10.1101/323030

**Authors:** Radoslaw Bomba, Witek Kwiatkowski, Roland Riek, Jason Greenwald

## Abstract

Interactions between biological membranes and disease-associated amyloids are well documented and their prevalence suggests that an inherent affinity exists between these two distinct molecular assemblies. Within the framework of our research project on amyloids and the origins of life, we hypothesized here that such interactions could increase both the sequence and structure space of peptide amyloids in a heterogeneous system and that if cooperative in nature, the interaction could be advantageous to the propagation of these entities in a prebiotic context. Thus, we have investigated the interplay between vesicle-forming fatty acids and amyloidogenic peptides, the respective precursors of lipids and proteins. Individually they are able to form ordered structures under a limited range of conditions with the bilayer of fatty acid vesicles and the cross-β core of amyloids both being repetitive structures that could in principle support a cooperative interaction. Here we report that an 8-residue basic peptide that can form an amphipathic β-strand, that is soluble at neutral pH and that can form amyloids above its pI at pH 11, is also able to cooperatively form novel co-aggregates of diverse structure with and in the context of simple fatty acids at neutral pH. Below the critical vesicle concentration (CVC) the mixtures of fatty acid and peptide yield a flocculent precipitate with an underlying β-structure. Above the CVC, the mixtures yield ribbon or tube-like structures that bear some of the hallmarks of amyloids yet have associated with them a significant amount of fatty acids, likely in a bilayer structure. In the context of the origin of cellular life these results expand the phase space of both peptides and fatty acids while providing a simple yet robust physical connection between two distinct biological entities relevant for life.

## 1. Introduction

Historically, the “origin of life” field has been polarized by competing, and seemingly exclusive hypotheses such as replication first (1), metabolism first (2) and compartmentalization first (3), or RNA world (4), peptide world (5, 6) and amyloid world (7–9). It is evident that these hypotheses tend to oversimplify the problem and that at one point in the origin or evolution of early life, the primary molecules of biology, namely nucleic acids, amino acids, aliphatic compounds and sugars became intimately intertwined. The emergence of these distinct molecular systems could have been sequential or separated in space or all at once as it has been postulated recently (10, 11). Physical interactions between the distinct systems may have stabilized them, strengthened their replication potential and/or enlarged their chemical and structural space in a way that would not be possible with a single type of biomolecule. Such mutually beneficial effects have been shown for DNA and peptide amyloids, for which DNA stabilizes the amyloid fibers and the amyloid supports DNA hybridization (12). Furthermore, the encapsulation inside of vesicles by amphiphilic membrane-like structures may be beneficial to the encapsulated system by offering protection from hydrolysis and by preventing loss of the system through diffusion, as has been demonstrated for example by the chemical replication of RNA inside fatty acid vesicles (13) or by protein translation within vesicles that contain the 80 different macromolecular species required for translation (14).

It is well-known that there is an inherent association of many disease-related amyloidogenic peptides with lipid bilayer membranes (15–18). Such interactions may arise from the fact that both can form periodic structures at the sub-nm level, thereby enhancing the potential for cooperative interactions. Within the framework of our investigations on the origin of life we hypothesized that fatty acids and amyloidogenic peptides, both arguably prebiotic entities, could also have cooperative interactions that stabilize each other’s structures and enhance their potential to undergo Darwinian selection. The sequence space of amyloidogenic peptides and the structure space of their fibers as well as the structure and phase space of fatty acids could be expanded on by cooperative interactions between peptides and fatty acids. Interactions of this sort in a simple system are demonstrated in the following results.

## 2. Material and Methods

### Materials

Sodium decanoate (TCI), sodium dodecanoate (Sigma-Aldrich), nonanoic acid (Acros organics), decanol (Sigma-Aldrich), Hydrochloric acid (Merck), sodium hydroxide (VWR Chemicals), potassium phosphate monobasic (Acros organics), sodium phosphate dibasic dodecahydrate (Acros organics), sodium metaborate (Sigma-Aldrich), acetonitrile HPLC Gradient grade (Fisher Chemical) and trifluoroacetic acid (Fisher chemical) were used without further purification.

The peptide (OV)_4_(GLS China) was synthesized using standard Fmoc chemistry on a Rink amide resin. The crude peptide was purified by reverse-phase HPLC on a Kinetex C18 5 μm 100 Å 250×10 mm column (Phenomenex) in a gradient of acetonitrile in 0.1% TFA. The purified peptide was quantitated by its calculated extinction coefficient at 214 nm of 6797 M^-1^cm^-1^ (19). Pure peptide was stored in lyophilized aliquots and dissolved in water just before use.

### Fatty acid solutions

Decanoate*-*decanoic acid (DA) and dodecanoate-dodecanoic acid (DDA) solutions were prepared by dissolving the corresponding sodium salts in phosphate buffer at room temperature or at 50 °C for decanoate or dodecanoate, respectively, and adjusting the pH of each solution to the desired pH with 1 M HCl, using a SevenCompact Mettler Toledo pH-meter fitted with a Hamilton BioTrode. After each addition of the pH titrant, the solutions were vortexed for 30 seconds to allow the solution to equilibrate before continuing the titration. After titration, the volume of the solution was adjusted to the desired concentration of fatty acid while keeping the concentration of phosphate constant at 50 mM. Nonanoate-nonanoic acid (NA) solutions were prepared at room temperature by mixing nonanoic acid in molar ratio 1:1 with sodium hydroxide and a volume of 0.5 M phosphate buffer equal to 1/10 of the final solution volume. Titration with HCl was carried out as for DA and DDA solutions. To prepare DA:decanol mixture, decanol was added to a sodium decanoate solution and vortexed for 1 minute. Subsequently titration and volume adjustment were done as for the other solutions.

### Formation of peptide-fatty acid mixtures

Directly before use, an aliquot of lyophilized (OV)_4_peptide was dissolved in water at 5 mg/ml and this was added to a fatty acid solution in a 1:9 volume ratio to give a final concentration of 0.5 mg/ml (OV) _4_. After peptide addition, the solutions were vortexed for 30 seconds and then incubated overnight at room temperature for NA and DA solutions, or at 50°C for DDA solutions.

### Circular dichroism (CD)

Peptide-fatty acid mixtures were analyzed by circular dichroism spectroscopy on a Jasco J-815 in a 0.1 mm quartz cell. The spectra were recorded from 260-190 nm at 50 nm/min, 2 nm band-pass, with a 2 s integration and averaged over 5 repetitions. Measurements were done at room temperature for NA and DA solutions and at 50 °C for DDA solutions.

### Critical micelle and critical vesicle concentrations

We employed four different methods to estimate the fatty acid critical micelle concentration (CMC) and critical vesicle concentration (CVC) under the conditions used in this study. The results of these measurements are in the Supporting Material and are summarized in Table 1.

**Table 1:** Measured CMC and CVC ranges for NA, DA and DDA solutions at pH 7.8

| Fatty acid | CMC |  | CVC |  |  | Literature |  |
| --- | --- | --- | --- | --- | --- | --- | --- |
|  | Fluorescence | DLS<sup>1</sup> | NMR | Turbidity | Fluorescence | CMC<sup>2</sup> | CVC<sup>2</sup> |
| NA | 50–70 mM | 50–70 mM | ND<sup>3</sup> | ND<sup>3</sup> | 150–200 mM | 110 mM | 85 mM |
| DA | 20–22 mM | 22–30 mM | 49–52 mM | 55–60 mM | 55–60 mM | 28.8 mM | 43 mM |
| DDA | 3.6–4.5 mM | NM<sup>4</sup> | 4.7–5.6 mM | 5 mM | 5–6 mM | 1.58 mM | 22.5 mM |

<sup>1</sup> CMC determined by the concentration at which signal began to fluctuate widely.
<sup>2</sup> CMC and CVC values from literature were determined using different procedures and at different pH (26, 27)
<sup>3</sup> No transition detected with this technique.
<sup>4</sup> DDA solutions not measured by DLS due to issues with solidification during samples preparation.

i. The fluorescence intensity of hydrophobic dyes as well as their fluorescence polarization can be used to measure the phase transitions of amphiphiles (20, 21). We chose 1,6-diphenyl-1,3,5-hexatriene (DPH) as a reporter molecule because both its fluorescence intensity and polarization are sensitive to amphiphile aggregation (22). DPH is minimally soluble in water and even at 5 µM remains mostly in aggregates that have low fluorescence intensity but a relatively high anisotropy. Above the CMC, micelles of the amphiphiles can solubilize some of the DPH leading to a moderate increase of intensity with a concomitant decrease in the fluorescence anisotropy. However, we found that DPH is significantly more soluble in vesicles than in micelles such that above the CVC, the fluorescence increases much more quickly with fatty acid concentration and there is another concomitant decrease in anisotropy. This fluorescence anisotropy CMC determination method was also tested with amphiphiles with known CMC values and the results are in good agreement with the literature values.
ii. Light scattering has also been shown to correlate with amphiphile aggregation (23) however, the CMC at pH 7.8 could not be precisely determined via light scattering as at this pH the signals became very unstable when the concentration approached the CMC. Each sample was filtered with 0.02 μm Anotop 10 membrane filter (Whatman). Measurements were performed on a Wyatt Dyna-Pro dynamic light scattering instrument. Only the scattered intensity and not the correlation function was used in the analysis.
iii. The onset of turbidity, measured as absorbance at 400 nm, was used to evaluate the CVC of the fatty acid solutions (24). The measurements were performed on a Jasco V-650 with a 1 cm path length and at room temperature for NA and DA and at 50°C for DDA solutions.
iv. The change in the ^1^H-NMR signal intensity of the methylene groups of fatty acids (NA: δ_H_ = 1.16 ppm, DA: δ_H_ = 1.20 ppm, DDA: δ_H_= 1.40 ppm) was employed to find the transition between micellar and vesicular state of amphiphiles. Below CVC the ^1^H signals are directly proportional to the fatty acid concentration. Above the CVC, the ^1^H signal intensity decreases with increasing concentration due to the large correlation time of the very large vesicles yielding fast spin relaxation. The CMC is not detected via this signal intensity method, most likely due to a fast exchange between monomers and micelles and a rather short correlation time of the fatty acids in the micelles and thus little change to their spin relaxation rate compared to monomers. Spectra were measured with Bruker Avance III 700 MHz and Bruker Avance III HD 600 MHz spectrometers, both equipped with triple resonance TCI CryoProbes. Samples were in 10% (v/v) D_2_O at 24.85 °C for NA and DA and at 50 °C for DDA. Data were processed and analyzed by Topspin 3.5 (Bruker).

All signal versus concentration data were analyzed using R software. Critical concentrations were determined from the NMR intensity data by fitting the linear portion of the trends and using the break from linearity as the CVC. For the fluorescence, light scattering, and turbidity data the CMC and CVC are reported as a range between the data points at which the trend in the data changed.

### Fluorescence anisotropy

The fluorescence anisotropy of 5 µM DPH in various fatty acid solutions was measured on a ISS K2 spectrofluorometer with 1 cm path length and excitation/emission wavelengths of 330/480 nm respectively. The samples were prepared by diluting 5 µl of a 1 mM DPH solution in acetonitrile into 1 ml of the sample to be analyzed and vortexing for 15 minutes. NA and DA samples were prepared and measured at room temperature while DDA samples were kept at 50 °C. The G-factor was measured independently for each sample and the recorded anisotropy values are the average of 10 measurements.

### Cryo-EM

Solutions to be analyzed were applied to a Quantifoil R2/2 Cu 300 grid and bloated for 10 s with bloat force 20. The grids were flash frozen in liquid ethane using a FEI Vitrobot and the images recorded with FC20 cryo FEI Tecnai electron microscope.

### Attenuated total reflectance Fourier transform infrared spectroscopy (ATR-FTIR)

Samples were centrifuged at 25k g for 5 min, pellets resuspended in 4 µl of supernatant and placed on the diamond ATR cell of Bruker Alpha-p ATR-FTIR spectrometer. Before the measurements samples were air-dried at room temperature. Fatty acid solutions without peptide were also measured as a control. An empty cell (air) was used as a blank spectrum.

### X-ray diffraction

Samples were centrifuged at 130k g with a Beckman Coulter Airfuge Ultracentrifuge, and the pellets were transferred into 0.3 mm glass capillaries, which were then flame-sealed on both ends to prevent drying. Scattering data were recorded with a Rigaku MicroMax-007HF source on a Mar345 image plate detector at room temperature with a 1 hour exposure. The scattering from an empty capillary was also measured and used for background subtraction. The diffraction patterns were radially integrated with FIT2D software (25) t? yield a plot of the scattered intensity as a function of d-spacing.

### HPLC analysis

In order to quantify the amount of peptide and fatty acid in the distinct aggregates, 100 µl samples containing aggregates (either flocculent or fibrillar) of (OV)_4_and fatty acids were centrifuged in a tared tube. The mass of the pellets were determined and then dissolved in 500 µl of 8 M Guanidine solution. Supernatants were analyzed without dilutions for NA and DA and the DDA supernatant was diluted 2-fold with 5% acetonitrile in H _2_O. Reversed phase analyses were performed on a Kinetex C8 5 μm 50 mm x 4.6 mm column (Phenomenex) connected to an Agilent 1200 HPLC system with an autosampler and diode array detector. The (OV)_4_peptide and fatty acid were resolved by a linear gradient of acetonitrile with 0.1% TFA (5-98%) at 3 ml/min. The injected amount of (OV)_4_was quantified based on its peak area and calculated extinction coefficient at 214 nm (19) of 6797 M^-1^cm^-1^ and fatty acids were quantified based on a calibration with known quantities of material. The pellet mass was mostly due to water (dried pellets had negligible mass) so the volume of the pellet was calculated from the mass of the wet pellet. This volume was then used to correct for the amount of material in the pellet fraction that is due to the soluble material trapped in the pellet.

## 3.Results and Discussion

To investigate the interactions between prebiotic peptides and fatty acids with a focus on a cooperative amyloid fibril and membrane bilayer formation, we designed the amyloidogenic peptide OVOVOVOV-NH_2_(subsequently referred to as (OV)_4_), with O for ornithine. It was predicted that the positive charge of (OV)_4_at neutral pH would prevent aggregation while allowing for favorable electrostatic interactions with the carboxylate of fatty acids. Indeed, at pH 7.8 the peptide is soluble up to 10 mg/ml and at 1 mg/ml adopts a random-coil-like conformation as evidenced by its CD spectrum (Fig. S1). We characterized the interaction of (OV)_4_with three simple vesicle-forming fatty acids: nonanoic (NA), decanoic (DA) and dodecanoic (DDA) acid. These fatty acids form vesicles in water at a pH near their pKa (26) but remain monomeric and micellar at a pH much above their pKa. It was therefore expected that at a pH above the fatty acid pKa, any cooperative effect between the peptide and fatty acid aggregation would be more pronounced and so a pH of 7.8 was chosen because it is higher than the pKa of all three acids (7.0, 7.3, and 7.5 for NA, DA, and DDA, respectively). Since both their critical micellar concentration (CMC) as well as the critical vesicle concentration (CVC) may be sensitive to pH, buffer strength and solution composition, we carefully measured these values for each fatty acid at the working pH of 7.8 (Table 1 and Figs. S2-S5). It is evident from the values in Table 1 that under the conditions used here (phosphate buffer, pH > pKa) the CMC and CVC values deviate from reported values measured at their respective pK_a_s. While this deviation is consistent with the known pH dependence of the fatty acid CVC, we nonetheless measured the CVC of DA at two different pH values (7.4 and 7.8) and found that in more acidic conditions, the CVC is significantly lower (25-28 mM) than for higher pH (49-52 mM). Additionally, at high enough pH fatty acids are not anymore able to form vesicular assemblies. Decanoic acid for example does not form vesicles above pH 8.2 (Fig. S4*D*).

Next, we prepared a concentration series of each of the fatty acids mixed with 575 µM (0.5 mg/ml) (OV)_4_at a pH of 7.8. In many samples, there was an immediate formation of a precipitate upon mixing of peptide and fatty acid and at the higher fatty acid concentrations, this precipitate appeared to go back into solution over a period of minutes to hours. This phenomenon was observed for all of the fatty acids, and the CD spectra of these mixtures revealed two concentration-dependent transitions separating three distinct states of the peptide (Figs. 1 and S6). At low fatty acid concentration, the peptide is in a random coil-like state but at a concentration of fatty acid close to the measured CVC (Table 1), the peptide precipitates and there is no CD signal. At a concentration above the measured CVC (Table 1), the CD spectra indicate that the peptide adopts a β-strand conformation (Figs. 1 and S6). The obvious conclusion of these data is that the transition concentrations observed in the CD spectra are very close to the CMC and CVC values for the fatty acids (illustrated in Fig. 2). Below the CMC, the peptide-fatty acid mixture is transparent and the CD spectra is indicative of a random coil-like conformation as it is in the absence of fatty acids. Near their CMC, fatty acids mixed with (OV)_4_produce a flocculent precipitate and the CD spectra of the mixtures are featureless, indicating that the peptide is contained in this flocculent precipitate. Above the CVC, the mixtures become translucent after an initial formation of precipitate and the CD spectra indicate that the peptide is now in a β-strand conformation. The loss of the CD signal in the concentration range between CMC and CVC is likely due to the large size of the flocculent aggregates that can no longer absorb light. To further characterize the mixtures, we centrifuged the translucent β-structured DA sample at 130k g and measured the CD spectrum of the soluble fraction, revealing the loss of the signal. Thus, the β-structured material in the translucent samples with fatty acid above its CVC is also an aggregate, but one that is small enough to absorb the UV light. The flocculent precipitate from the samples containing the fatty acid below its CVC that lacked a CD signal was further characterized by ATR-FTIR. The FTIR spectra had absorbance bands near 1625 cm^-1^, indicating that the peptide in the flocculent precipitate was also in a β-strand conformation (Fig. S7).

**Figure 1:**
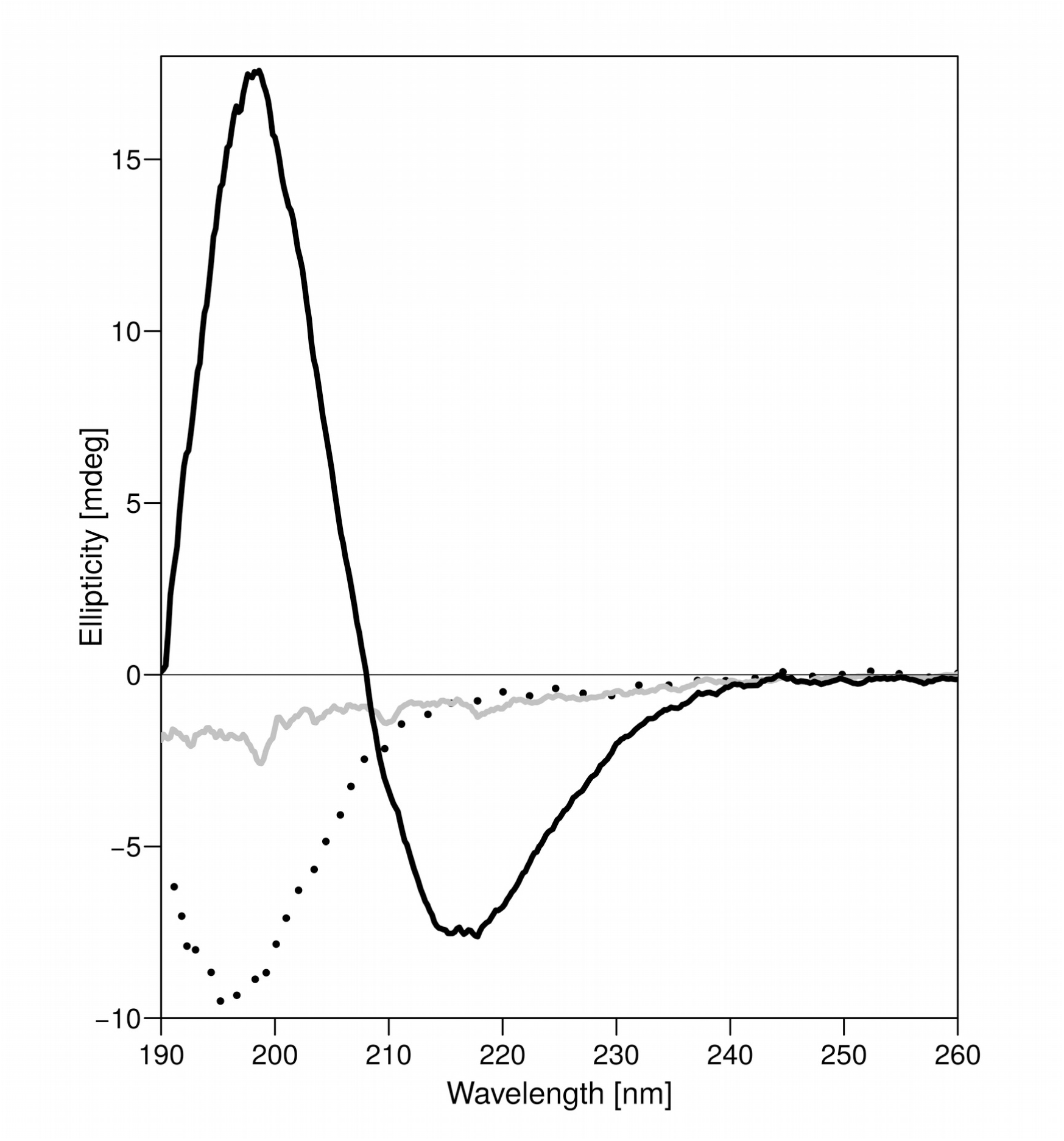
The concentration dependence of the CD spectra of DA solutions with (OV)_4_. The CD spectra of three solutions of 575µ M (OV)_4_ with either 1.8 mM DA (dotted line). 18 mM DA (solid grey line) or 72 mM DA (solid black line) are shown. At 1.8 mM DA the CD spectrum is random-coil-like, at 18 mM DA it is nearly featureless and at 72 mM it is β-sheet-like.

**Figure 2:**
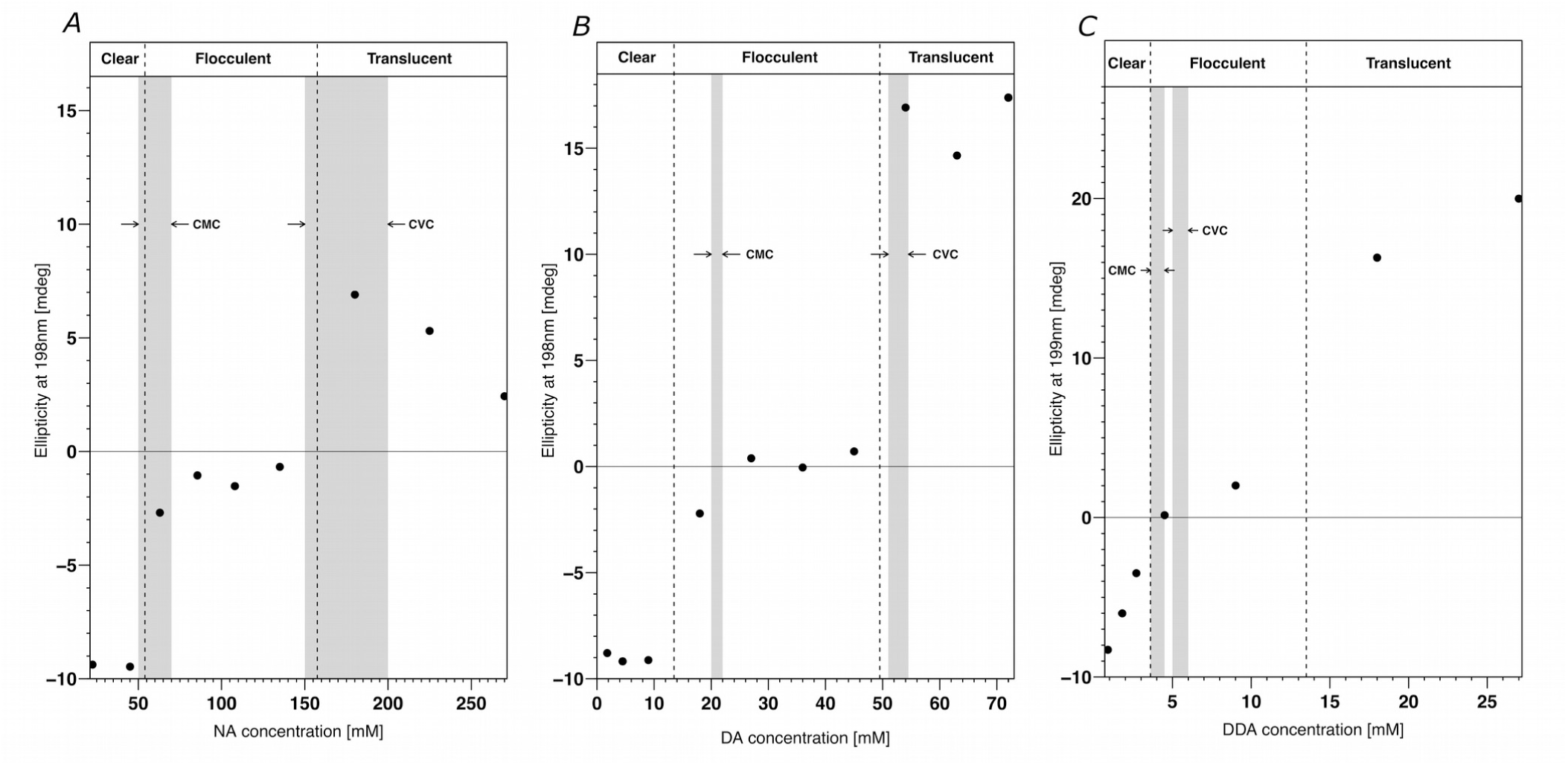
Correlation of (OV) _4_ secondary structure with fatty acid CMC and CVC. The β-strand secondary structure of the peptide in the fatty acid mixtures as monitored by its CD signal at 198 nm is plotted against the fatty acid concentration in solutions of NA (*A*), DA (*B*) and DDA (*C*). Negative values are associated with a random coil-like structure of the peptide and positive values with β-sheet secondary structure. The dashed lines roughly demarcate the boundaries between the visible changes in the mixtures from clear solution to flocculent precipitate to translucent solution with increasing concentration of fatty acid. The grey bands represent the ranges of CMC and CVC values for the fatty acids as measured independently in the conditions used in this study (see Figs. S2-S5) and indicate that the transitions of peptide-fatty acids mixtures occur at very similar concentration ranges as do the phase transitions of the pure fatty acids themselves. The deviation of the DDA mixtures from this trend can be explained by the fact that its concentration is low enough that the binding of DDA to the 575 μM (OV) _4_ consumes a significant amount of the fatty acid, effectively shifting the flocculent-translucent boundary.

This interesting correlation of the CMC/CVC values of the three fatty acids with the characteristics of the (OV) _4_-fatty acid mixtures suggests that mixtures of fatty acid micelles with (OV) _4_form the flocculent precipitate while fatty acid vesicles are required to form the translucent peptide precipitates. This correlation was further probed by modulating the CVC of DA by the addition of decanol (DOH) as it is known that the CVC of fatty acid solutions are lowered in mixtures that contain small amounts of the corresponding fatty alcohol (26). A 10:1 DA:DOH mixture has been reported to have a CVC of approximately 20 mM (26) in contrast to the CVC of 50 mM in absence of fatty alcohol (Table 1). Indeed, when (OV)_4_was mixed with a 39.6 mM (total amphiphiles) 10:1 DA:DOH solution, the translucent precipitate was formed and its CD spectrum contained the β-strand signal (Fig. 3), while at a concentration of 45 mM DA in the absence of fatty alcohol, a flocculent precipitate is formed and the CD spectrum of the mixture is nearly featureless (Fig. 3). This experiment further supports the correlation between CVC and the formation of a translucent precipitate.

**Figure 3:**
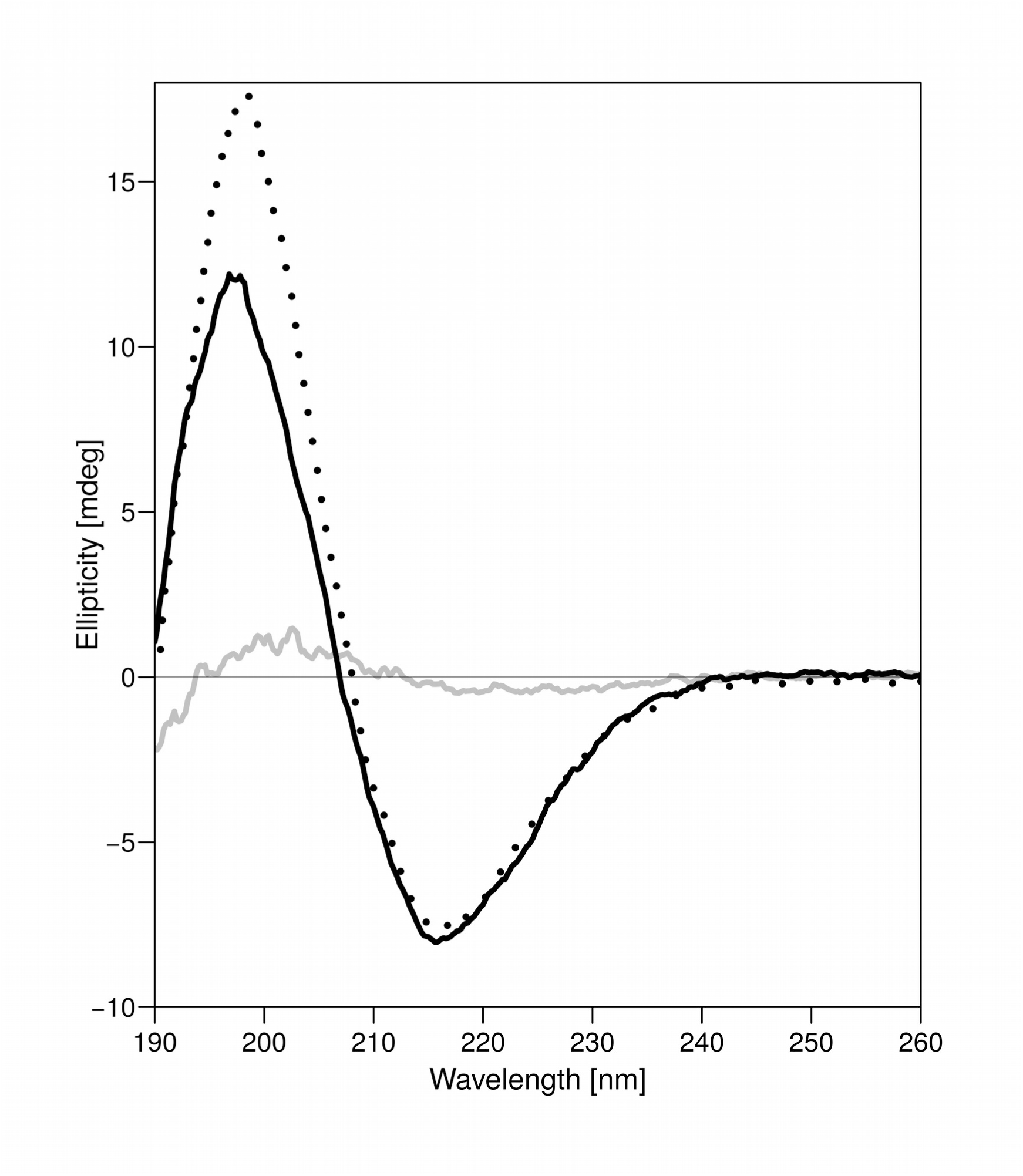
Fatty alcohol lowers both CVC and the concentration at which translucent precipitate is formed. Depicted are three CD spectra for 575 μM solutions of (OV) _4_ with either 72 mM DA (dotted line), 45 mM DA (solid grey line) or a 10:1 DA/DOH mixture with 39.6 mM total amphiphiles (solid black line). The β-sheet–like spectrum absent in the 45 mM DA solution due to the formation of the flocculent precipitate is present in the lower concentration DA solution with a 1/10 molar ratio of fatty alcohol. This amount of DOH decreases the CVC of the system from approximately 55 mM to approximately 20 mM.

We attempted to characterize the different peptide-fatty acid aggregation states and found that their formation is quickly reversible. Upon dilution of the mixtures with buffer, the precipitates dissolve and the diluted samples have random coil-like CD spectra. Therefore, for measuring the content of fatty acid and peptide in the aggregates, we could not simply rinse the aggregates but rather had to subtract the amount of soluble peptide and fatty acid remaining in the wet pellet (see methods). This analysis showed that the flocculent and translucent aggregates do in fact contain both fatty acids and peptide. The molar ratios of fatty acid to peptide in the flocculent precipitates are approximately 3 for NA and DA and 9 for DDA while in the translucent precipitate, the ratio for NA and DA is approximately 7 (DDA translucent precipitates could not be assayed due to the low melting temperature of DDA that causes it to solidify during centrifugation) (Fig. S8). The doubling of the fatty acid content in the translucent precipitate compared to the flocculent precipitate would be consistent with a conversion from a monolayer to a bilayer structure. This is particularly noteworthy in light of the facts that a micelle to vesicle conversion is also a change from a monolayer to bilayer and that the formation of the two types of precipitates is correlated with the fatty acid CMC and CVC. The high fatty acid content that was measured in the DDA flocculent precipitate also provides an explanation for the appearance of the flocculent precipitate above the measured CVC (Fig. 2*C*). The CMC and CVC of DDA are so similar and so low that the binding of DDA to the 575 μM (OV)_4_consumes a significant amount of the fatty acid, thereby effectively shifting the flocculent-translucent boundary by about +5 mM (i.e. ∼9×575 μM) from the CVC. Such a shift may also be present for the other fatty acid mixtures however it would be less noticeable due to their relatively large CVC values.

In order to better characterize the mesoscopic structure of peptide-fatty acid mixtures, we analyzed the flocculent and translucent precipitates by cryo-EM. The DA-(OV)_4_flocculent precipitate was generally too dense to get much information from the EM images, however, in a few instances we could detect fibril-like structures, suggesting that the β-structure detected by FTIR is in fact an amyloid (Fig. 4*A*). The translucent precipitates of all the mixtures consisted primarily of straight ribbon-or tube-like structures. In the case of the DA-(OV) _4_mixture, these straight structures have a uniform width of about 25 nm and lengths of up to several microns (Fig. 4*B*). The translucent NA-(OV)_4_mixture was similar in nature, in line with its similar fatty acid content, however, instead of a uniform width there is a narrow distribution ranging from 20 to 30 nm. In contrast, the DDA-(OV)_4_mixtures gave structures with a much larger variation in width from 15 nm to 80 nm (Fig. 4 *D, E*). Amyloid fibrils are often found to be composed of bundles of protofilaments that can be either twisted, straight or displaying some other combination of quaternary structures (28). Such bundling is detected in the (OV)_4_fibrils formed without fatty acid at pH 11 (Fig. 4*F*). Also, the ribbon-like structures in the translucent precipitates often have a detectable striation that indicates that they may be composed of many smaller fibrils that have assembled into a larger ordered structure (Fig. 4*C*). Still, the structures in the DA-(OV)_4_translucent precipitate are unusual in the homogeneity of their width distribution. Flat ribbons made of side-to-side stacked protofilments would be expected to be found with various numbers of filaments and thus of varying widths, as is the case for the DDA-(OV)_4_translucent precipitates (Fig. 4*E*). Thus, a higher order geometrical restraint appears to be at play in the DA-(OV)_4_structures. This could, for example, arise from a curvature in the “ribbons” such that they form closed nanotubes as has been observed for other amyloids (29–31). This interpretation would also explain the complete absence of tilted or side-on views of the structures in any of the hundreds of structures that we visualized by cryo-EM.

**Figure 4:**
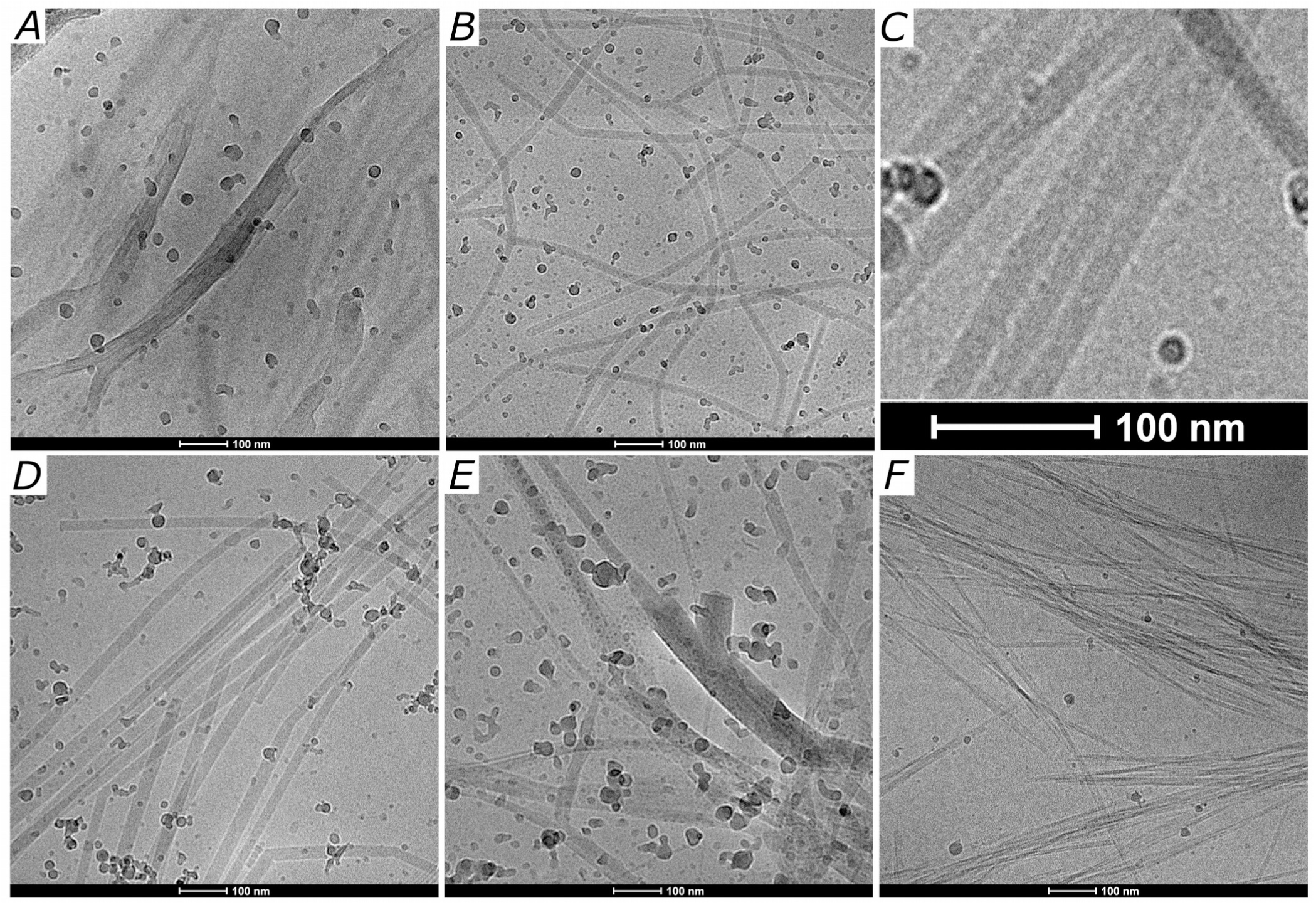
Morphology of fatty acid-(OV) _4_ precipitates compared to pure (OV)_4_ amyloids. Cryo-EM micrographs of the flocculent precipitate (*A*) and translucent precipitate (*B*) formed in mixtures of DA and (OV) _4_.The flocculent precipitate is usually too thick for observing detailed morphological features, but in a few instances like the one depicted in (*A*), fibrillar structures are visible. The translucent precipitate is characterized by straight fiber-like structures that have a uniform width of about 25 nm and lengths of up to several microns. In many of the tube-like structures, there is a visible striation indicating that they are composed of an alignment of smaller fibrils (*C*). The translucent precipitates of (OV) _4_ mixtures with NA (*D*) and DDA (*E*) also reveal fiber-like structures however they present a distribution of widths. The NA samples have a narrow distribution while the DDA/(OV) _4_ mixtures have structures with a much larger variation in width. The amyloid fibrils of (OV) _4_ that form at pH 11 (*F*) are characterized by many straight fibers that form bundles of varying numbers of proto-filaments.

The peptide-fatty acid precipitates were further characterized by X-ray diffraction which revealed some similarities between the flocculent and translucent precipitates that are also shared with the (OV) _4_pH 11 fibrils. The DA-(OV)_4_precipitates as well as the (OV)_4_pH 11 fibrils have a reflection at 4.7 Å which corresponds to the spacing between strands in a β-sheet. The other similarities include reflections near 7.8 Å and 11.7 Å (Fig. 5). The DA-(OV)_4_precipitates have as well an intense reflection near 24 Å that is mostly absent in the pH 11 fibrils of the peptide without DA. Unique to the fatty acid translucent precipitate is a broad reflection at 30 Å, whichmaybe indicativeofafatyacidbilayer. We were not able to detect any spontaneous alignment of the samples in the glass capillary and did not want to dry the samples to induce alignment for fear of altering the structures in the co-aggregated samples. Therefore, despite the presence of reflections typical of a cross-β-sheet at 4.7 Å and 8-10 Å, the x-ray diffraction alone does not provide information on the orientation of the β-sheets within the ribbon-like structures.

**Figure 5:**
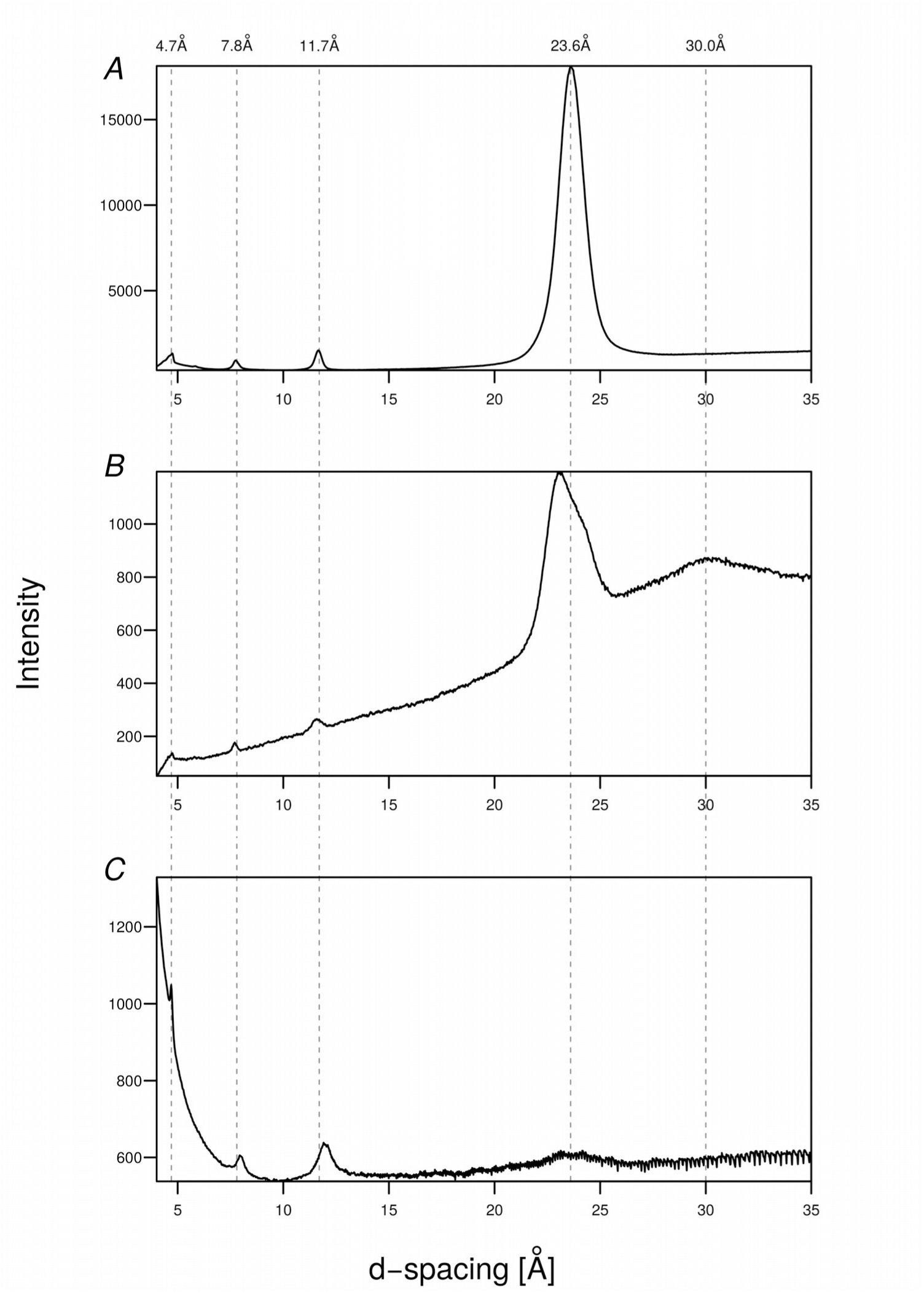
X-ray diffraction of fatty acid-(OV) _4_ precipitates compared to pure (OV) _4_ amyloids. The X-ray diffraction images of flocculent (*A*) and translucent (*B*) precipitates from DA-(OV) _4_ mixtures at 18 mM DA and 63 mM DA, respectively, are shown. In (*C*) the diffraction of (OV) _4_ amyloids formed at pH 11 in the absence of fatty acids is shown. All of the various aggregates have a reflection at 4.7 Å corresponding to the spacing between strands in a β-sheet. The other similarities include reflections near 7.8 Å and 11.7 Å. The DA/(OV) _4_ precipitates also have an intense reflection around 24 Å that is mostly absent in the pH 11 amyloid fibrils that lack DA. Unique to the translucent precipitate is a broad low resolution reflection at 30 Å which may be indicative of a bilayer. The diffraction patterns were radially integrated with the FIT2D software t? yield a plot of the scattered intensity as a function of d-spacing.

Taken together, our results suggest that the presence of vesicles is necessary to form the ribbon/tube-like aggregates with the peptide (OV)_4_. However, since we were unable to detect the presence of NA vesicles by NMR or turbidity up to its solubility limit of approximately 1 M at pH 7.8, it is also possible to view the peptide-fatty acid interaction as creating a separate phase whose boundaries, under the conditions used in this study, are very close to the phase boundaries of the fatty acids themselves. Likewise, the large changes in the fluorescence intensity and anisotropy of DPH between 150 and 200 mM NA could be due to a similarly cooperative interaction in which DPH induces bilayers of NA outside of the normal pH range for NA vesicles (Fig S3*A*). In any event, the observed correlation between CVC and (OV)_4_-fatty acid aggregate structure suggests that the translucent precipitate may have some aspects of a fatty acid bilayer. This is further supported by the doubling of the fatty acid content relative to peptide in the translucent precipitate compared to the flocculent precipitate as well as the X-ray reflection with a 30 Å d-spacing unique to the translucent precipitate. Thus, the fatty acid not only increases the pH range at which (OV)_4_ is able to aggregate, it also changes drastically the structures that are formed compared to peptide alone at alkaline pH. Conversely, the peptide changes the structure of the fatty acid aggregates and increases the pH range (at least for NA) at which aggregation occurs.

Viewed from a prebiotic perspective, the enhancements of both the structural space of these molecules as well as the environmental conditions under which they are capable of forming repetitive structures can be regarded as selective advantages. Considering the replicative properties of both amyloids and fatty acids (32, 33) cooperative interactions of the kind we report here may have led to new functions that could be beneficial to the propagation of either one or both of these molecules. These functions may for example include novel catalytic activities, as has been reported for numerous amyloidogenic peptides (34). In addition, the rapid and reversible soluble – aggregate nature of the peptide-fatty acid interactions may be a prebiotic-relevant unique functional property of the co-aggregates, since amyloids are usually rather stable (8) and reversibility is more often associated with the biologically functional amyloids (35, 36). The cooperative interaction that we describe here would be therefore one way to explain the original connection between peptides and fatty acids that eventually led to cellular life.

